# Taking a deeper look: Quantifying the differences in fish assemblages between shallow and mesophotic temperate rocky reefs

**DOI:** 10.1101/449959

**Authors:** Joel Williams, Alan Jordan, David Harasti, Peter Davies, Tim Ingleton

**Affiliations:** Fisheries Research, NSW Department of Primary Industries, Locked Bag 1, Nelson Bay, NSW 2315, Australia; New South Wales Office of Environment and Heritage, 59-61 Goulburn St, NSW, Sydney, 2001, Australia

## Abstract

The spatial distribution of a species assemblage is often determined by habitat and climate. In the marine environment, depth can become an important factor as degrading light leads to changes in the biological habitat structure. To date, much of the focus of ecological fish research has been based on reefs in less than 40 m with little research on the ecological role of mesophotic reefs. We deployed baited remote underwater stereo video systems (stereo-BRUVS) on temperate reefs in two depth categories: shallow (20-40m) and mesophotic (80-120m), off Port Stephens, Australia. Sites were selected using data collected by swath acoustic sounder to ensure stereo-BRUVS were deployed on reef. The sounder also provided rugosity, slope and relief data for each stereo-BRUVS deployment. Multivariate analysis indicates that there are significant differences in the fish assemblages between shallow and mesophotic reefs, primarily driven by *Ophthalmolepis lineolatus* and *Notolabrus gymnogenis* only occurring on shallow reefs and schooling species of fish that were unique to each depth category: *Atypichthys strigatus* on shallow reefs and *Centroberyx affinis* on mesophotic reefs. While shallow reefs had a greater species richness and abundance of fish when compared to mesophotic reefs, mesophotic reefs hosted the same species richness of fishery targeted species. *Chrysophrys auratus* (pink snapper) and *Nemodactylus douglassii* (grey morwong) are two highly targeted species in this region. While *C. auratus* was numerically more abundant on shallow reefs, mesophotic reefs provide habitat for larger fish. In comparison, *N. douglassii* were evenly distributed across all sites sampled. Generalized linear models revealed that depth and habitat type provided the most parsimonious model for predicting the distribution of *C. auratus,* while habitat type alone best predicted the distribution of *N. douglassii*. These results demonstrate the importance of mesophotic reefs to fishery targeted species and therefore have implications for informing the management of these fishery resources on shelf rocky reefs.

## Introduction

The spatial distribution of a species assemblage is strongly determined by habitat and physical conditions [1,2], and in the marine environment depth is an important factor [3–5]. On the inner continental shelf the decreased light conditions with increasing water depth results in a change from macroalgal to sessile invertebrate dominated habitat composition [6,7]. In temperate waters this change occurs at depths of around 20-30 m, although variations occur reflecting localised conditions [8–10]. To date, much of the research on rocky reefs on the inner shelf has been focussed on reefs in depths less than 20 m reflecting the widespread use of SCUBA to conduct such surveys. There are few standardised tools to quantitatively survey fish at greater depths. This is despite the significant range of pressures on deeper rocky reefs across the continental shelf from activities including commercial and recreational fishing that target reef associated species. In recent decades there is anecdotal evidence that recreational fishers have increase technical capacity such has side scan or multibeam sonar and electric reals and are therefore, targeting deeper reefs. Previous these reefs may have provided refuge for older mature individuals.

Mesophotic reefs are those characterised by middle to low levels of light [7,11–13]. The recent worldwide expansion of multibeam acoustic surveys of continental shelf waters has revealed that they form extensive areas of habitat in many regions [9,14,15]. Mesophotic reefs are often continuous with shallow reefs resulting in a strong connectivity across a large depth gradient, a feature common in the Great Barrier Reef Australia, north eastern Brazil and the Hawaiian Archipelago [6,7,16,17]. They can also form discontinuous areas that are interspersed among areas of unconsolidated habitat, such as in the Gulf of Carpentaria, Australia [12,18]. While the number of studies on mesophotic reef has increased significantly over the past decade [13,19], the majority of this research is focussed in the tropics where such reefs are usually dominated by corals [12,13]. Conversely, temperate mesophotic ecosystems tend to be dominated by sponges and octocorals [9,14]. In comparison to the tropical mesophotic coral ecosystems, very little is known about temperate mesophotic ecosystem, particularly the link between fish assemblages, habitat structure, and connectivity with shallower reefs.

Temperate mesophotic reefs have important biodiversity, social and economic values [11], so understanding the characteristics of their associated fish assemblages is fundamental to effectively managing them. Habitat type (coral, sponge, bare) and complexity (relief, rugosity, curvature) are known to be important in structuring fish assemblages [20–25]. Habitat complexity is considered as the variance in surface structure of the reef and can be defined in terms of relief, slope, rugosity, surface area, and other factors [22,26]. The link between habitat complexity and fish assemblages has been well researched, with many studies showing positive relationships between complexity and fish abundance, biomass and diversity [21,27–32].

As mesophotic reefs often occur adjacent to inshore shallow reefs, some connectivity across the depth gradient might be expected. It was hypothesised in the late 1990’s and early 2000s, for example, that mesophotic reefs provide refuge for some fish species [33–35]. This hypothesis assumed that mesophotic reefs were isolated from most of the stressors that impact inshore shallow reefs such as coral bleaching, pollution, habitat loss and some forms of fishing [35]. For temperate reef systems, there are insufficient data over sufficient temporal scale to make generalised conclusions about the extent or nature of any habitat connectivity between shallow and deep components. There is empirical evidence based on genetics and observations that connectivity between mesophotic and shallow reefs occurs, and this has been observed for over-exploited fishery target species[19,35]. On the other hand, there is also some evidence that mesophotic reefs are not merely extensions of shallow reefs, but host a unique species assemblage and would benefit from increased conservation management [3,4,36].

Surveys of mesophotic reefs have historically been logistically difficult and expensive due to the need for large offshore vessels and the lack of detailed information on their distribution and structure [23,37,38]. Earlier studies used coarse scale maps generated through the aggregation of information from commercial fishers, historical hydrographic data and targeted single beam acoustic surveys [39–41]. More recently, the expansion of swath acoustic surveys in has resulted in high resolution maps of continental shelf rocky reefs based on the interpretation of bathymetry and backscatter [42]. The recent development of cost effective and easy to deploy underwater video equipment has also meant a move away from destructive survey methods such as gillnets, droplines and traps, which are also often not suitable for use in sensitive or protected areas. Baited remote underwater video (BRUV) is now commonly used to survey fish assemblages, and advances in camera housings, lights and study designs have enabled the deployment of cameras onto deeper habitats enabling non-destructive sampling of fishes across continental shelf waters[30,43–45].

While there has been some assessment of fish assemblages on shallow reefs in temperate eastern Australia [46–49], there has been no comparable assessment done on mesophotic reefs. The lack of knowledge on fish assemblages and habitat composition of mesophotic reefs in this region increases the uncertainties associated with marine spatial planning and evaluation of management effectiveness. This is particularly important in the case of marine protected areas which usually have specific objectives about conserving the biological diversity of a representative range of habitats and associated assemblages. Our study focussed on an area of the inner continental shelf within the Port Stephens-Great Lakes Marine Park (PSGLMP) and the adjacent Hunter Marine Park (HMP). The PSGLMP extends from the tidal limit to the 3 nm extent of State coastal waters, with the HMP extending from this boundary to 25 nm offshore. Specifically, the aim of our study was to quantify and compare the spatial distribution of fish assemblages on shallow (20-40 m) and mesophotic (80-110 m) temperate rocky reefs, and relate these to habitat composition. This study will extend knowledge of temperate mesophotic reefs generally, but will also inform future decision-making in these two marine parks to improve effective management of these reefs and to design better targeted monitoring programs.

## Material and methods

### Study site

This study took place along an ∼40 km length of coastline between Port Stephens and Seal Rocks within the waters of the PSGLMP and HMP in New South Wales, Australia (−32°S; Fig. 1). This region is dominated by temperate species, although some tropical vagrants often arrive in summer, reflecting the strong presence of the East Australian Current that originates in the tropics [50–53]. This study was conducted during the austral spring of 2016, a period of cool water influence in this region when surface temperatures are around 15-17°C.

**Fig. 1.**
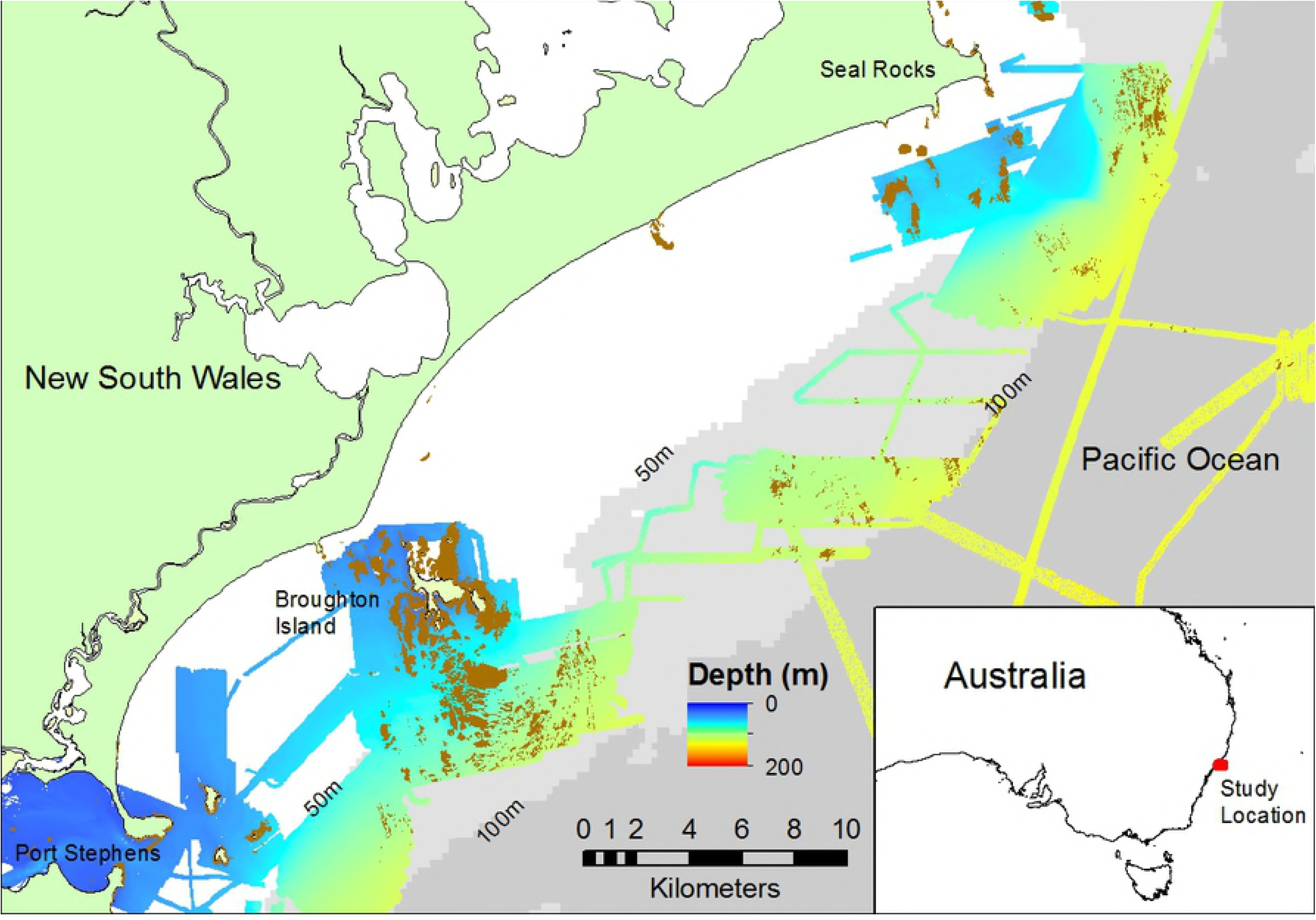
Map of the study area from Port Stephens to Seal Rocks on the east coast of Australia. The area that has been mapped using a swath acoustic sounder is indicated by the rainbow shaded area, while the 50m and 100m contours are represented by a differing shade of grey (source: GeoScience Australia). The brown shapes represent hand digitised reef with profile. Shallow stereo-BRUV deployments are delineated by the green circles and mesophotic stereo-BRUVs deployments are delineated by the blue circlers. Circles with a cross are BRUV deployments within a no-take area. Inset: Study location on the east coast of Australia.

Appropriate ethics (NSW Department of Primary Industries: ACEC REF 10_09) and fieldwork (NSW Department of Primary Industries research activities in NSW state waters: Permit No. P01/0059(A)-4.0 and research activities inside a marine park: Permit No. PSGLMP 2018/010) permits were obtained for this work.

### Reef mapping

In order to evaluate the extent, distribution and structure of rocky reefs in the study region, swath acoustic bathymetry and derived habitat data were collated from previous surveys [9]. In addition, targeted acoustic surveys were conducted in the region [54].

All acoustic surveys were conducted using a 125 kHz Geoswath interferometric swath system. Position and vessel motion for sonar acquisition was provided using a POS MV (Applanix, Canada) with Real-time Kinematic height and positional Virtual Reference Station corrections through SmartNet across the Telstra Mobile 3G network using Hypack (Hypack USA) acquisition software. Real-time ephemeris data were saved in POSPac log files for post-processing and calculating a 3 min forward-backward smooth for improved SBET in Single Base Station Mode. Mean SBET positional accuracies were improved to be better than 0.1 for X, Y and Z at nadir. SBETs were applied to Geoswath data before rough processing using amplitude, box, across-track and along-track filters in GS+. Data were exported as GSF for further data cleaning and cube modelling of soundings and production of a final digital elevation model. Backscatter data were output from GS+ in XTF data and then mosaiced using the Sidescan Solo module within Fledermaus FMGT. Reef extent was hand digitised from hillshaded bathymetry and derived slope layers and identified as ‘reef with profile’.

The Spatial Analyst tool and Benthic Terrain Modeller add-on in ArcGis v10.3.1 were used to analyse the cleaned bathymetric data. Fifty m and 100 m radius buffers around individual BRUVS were used to calculate the mean, standard deviation and range for relief, rugosity, ruggedness, curvature and slope at the specific sites where fish assemblages were quantified. Due to the 200 m separation between BRUVS, this ensured there were no overlaps. Pearson’s correlations were used to assess data obtained from the 50 m and 100 m radii for correlation.

### Sampling fish assemblage

Stereo baited remote underwater video (stereo-BRUV) was used to sample the fish assemblages at two depth strata, shallow reef (20-40m) and mesophotic reef (80-110m). Sampling sites were chosen using randomly selected grid references and a 1×1 km grid overlay on the plotted swath acoustic bathymetry maps. Each site consisted of four replicate stereo-BRUV deployments that were selected using 200×200 m grids to ensure each replicate was randomly selected yet spatially independent. Sites were located within Port Stephens - Great Lakes Marine Park (PSGLMP) and the Hunter Marine Park (HMP), with 48 BRUV deployments located within no-take areas within the PSGLMP, and 59 BRUV deployments in areas that are fished. Hence, fishing status was included as a factor in the modelling (see Table 1).

**Table 1.**
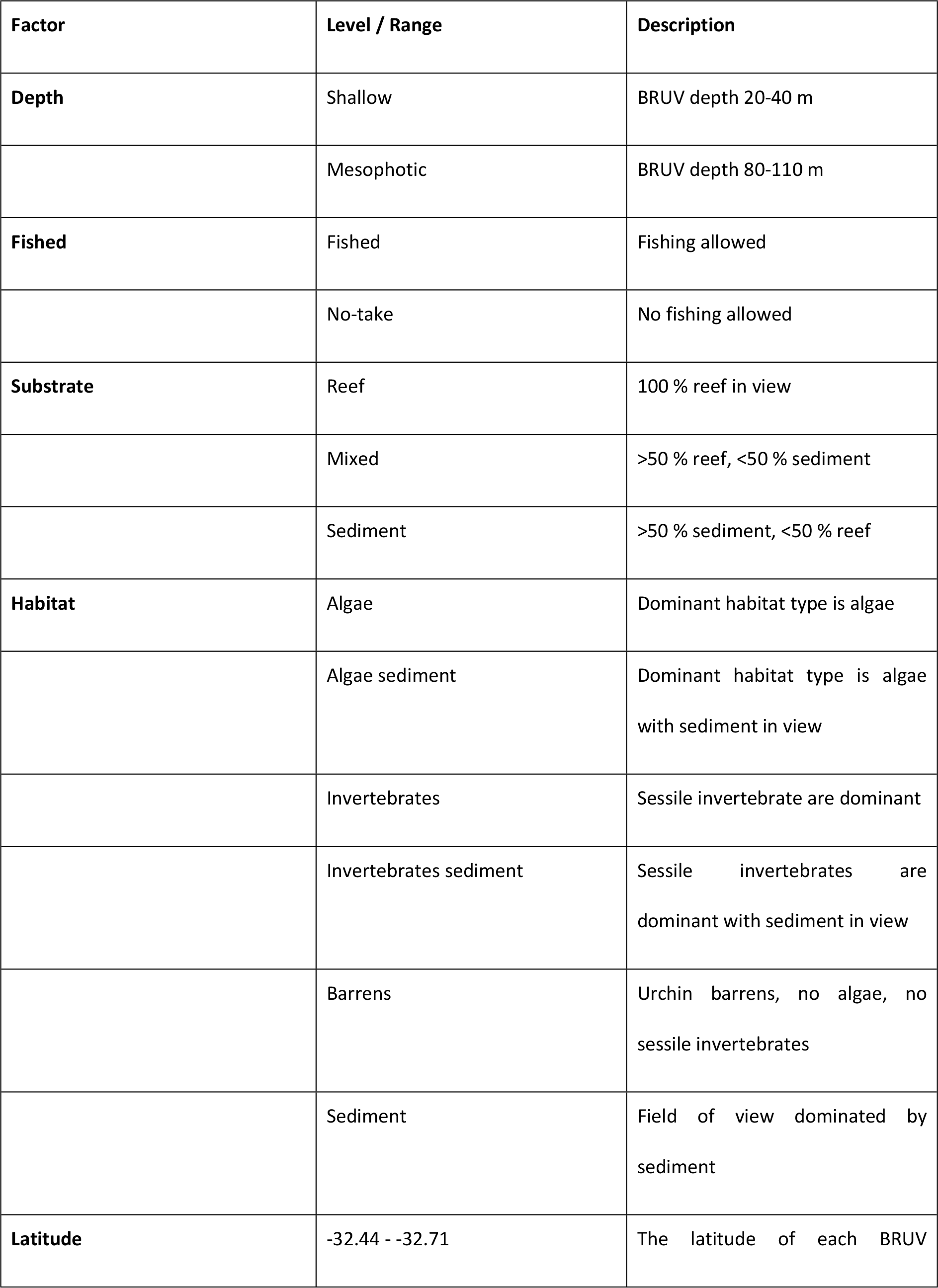
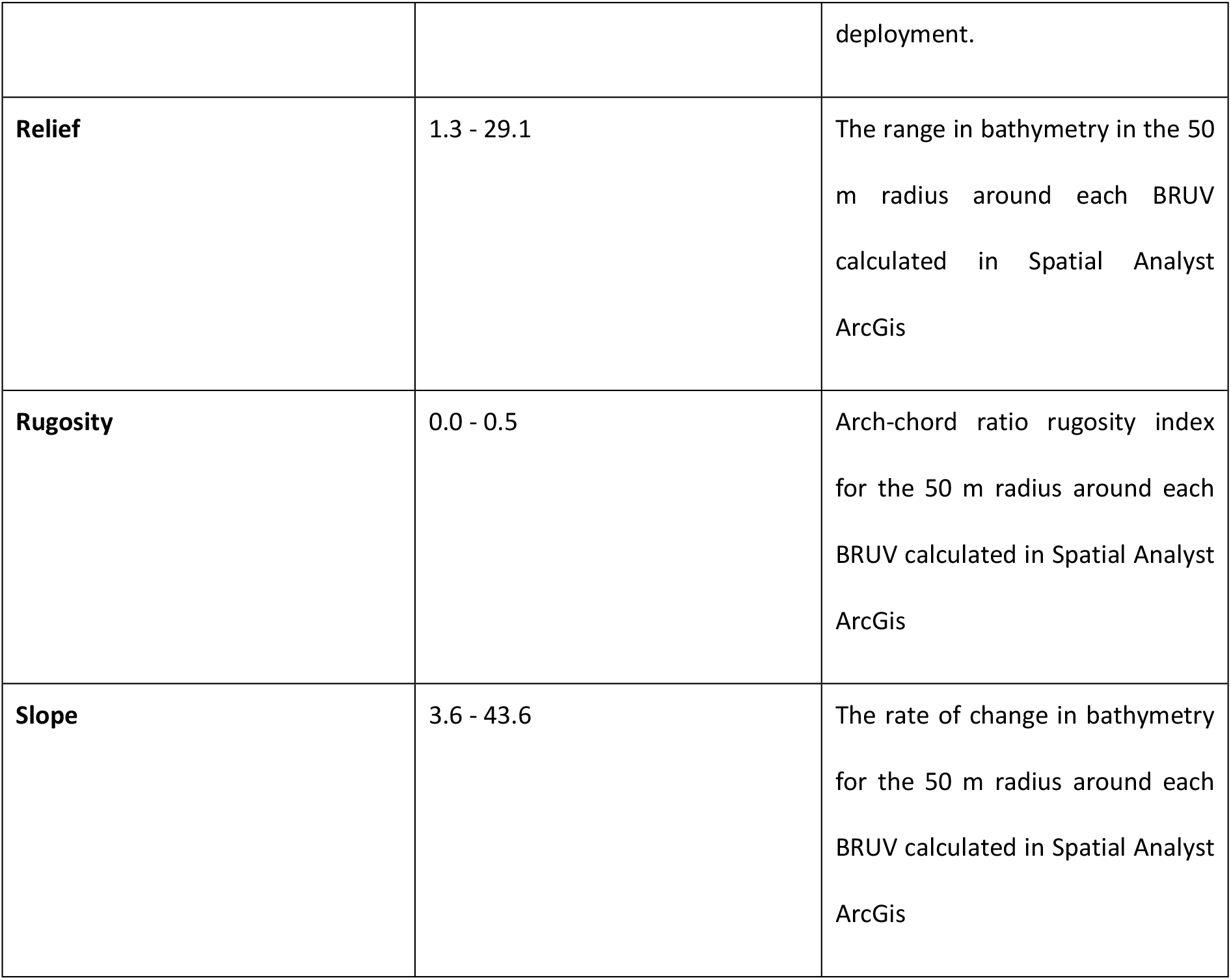
Description of factors used in both GAMM and RDA modelling.

Each deployment targeted rocky reef, and a deployment was considered successful if the stereo-BRUV landed on or immediately adjacent to rocky reef and when both the reef/benthos and water column could be viewed clearly. If a replicate was located over soft sediment it was moved to the nearest area of reef. Stereo-BRUV’s were deployed for a period of 30 minutes which has been determined to be a sufficient time to obtain a representative sample of the fish community [55].

Each BRUV unit consisted of two Canon HG25 video cameras with a wide angle lens that were housed in two custom made SeaGIS Lty Ltd housings (http://www.seagis.com.au). Approximately one kilogram of pilchard (*Sardinops sp.*) was crushed in a plastic mesh bait bag and attached to the BRUV frame at the end of a 1.5 m long PVC pole. Due to the low light levels at depths >80m, we used Raytech subsea lights mounted to the centre of the BRUV frame at sites below that depth. A blue light was used as it is likely that the 450-465 nm wavelength is below the spectral sensitivity range of many fish species and therefore will have minimal effect on the fish behaviour [56]. On several occasions, white light was used to confirm identifications of fish species and to collect qualitative data on habitat type.

Video imagery collected by BRUVs was scored using standard metrics including scoring relative abundance (MaxN) as the maximum number of fish occurring in any one frame for each species. MaxN is widely accepted as the best method for estimating relative abundance from video footage [43]. All fish were identified to the lowest taxonomic level possible, ideally to species level. For each stereo-BRUV deployment, the length of each *Chrysophrys auratus* and *Nemadactylus douglasii* observed at the time of MaxN was measured as total length (tip of fish nose to tip of tail fin). These two species are considered fishery target species and are often used as indicator species in BRUV surveys [55]. All BRUV video analysis and scoring was done using the EventMeasure software (www.seagis.com). The video footage was also used to categorise substrate type and habitat type as factors in an attempt to relate species and species assemblage data to the environment and habitat. (Table 1).

### Data analysis

To examine patterns in species assemblages, we used redundancy analysis (RDA). RDA is related to principal components analyses (PCA) and is based on Euclidean distance, implying that each species is an on axis orthogonal to all other species, and sites are points in this multidimensional space [57]. Due to a number of schooling species occurring in high abundances, all species were Hellinger transformed before doing a forward stepwise model selection using a suite of explanatory factors (Table 1) to select the factors that best explained the dissimilarity in the species assemblage. The function “ordiR2step” from the “Vegan” package in R was used to select the most parsimonious model [58]. Permutation tests were used to test for the statistical significance of each marginal term. A triplot was used to visually determine and display the strength of the relationships between species assemblage and the explanatory factors that underpin the variation in species assemblage between BRUV deployments.

To investigate the spatial distribution of the fish assemblage across shallow and mesophotic reefs we used generalised additive mixed models (GAMMs). These can incorporate the non-linear patterns and overdispersion often encountered with spatially structured ecological studies. A suite of response variables was chosen *a priori* and these included species richness, total relative abundance, the most speciose families (Labridae, Monocanthidae and Carangidae) and species that are either abundant (silver trevally *Pseudocaranx georgianus* and velvet leatherjacket *Meuschenia scaber*) or of fishery interest (*C. auratus*, *N. douglasii*,). We also modelled the relative abundance of all recreationally and commercially targeted species pooled together. Recreationally targeted species were determined from [59], while commercially targeted species were selected from the assessment reports for the ocean trawl and trap & line fishery assessment reports[60]. Site, a cluster of four BRUV deployments, was used as the random factor.

Prior to any modelling, all data were explored using scatter and boxplots to assess for correlations between covariates and outliers in the response variables. To minimise the risk of overfitting any models, if two covariates had a Pearson’s correlation >0.7, then the variable that made the least ecological sense in explaining the distribution of fish was removed.

A forward stepwise method was used to select the ‘best’ model based on AIC. The first step ran models with individual predictor variables and the model with the lowest AIC was then selected. Step two ran models including the first predictor variable with all other variables and again selected the variable with the lowest AIC. This was repeated until the difference in the AIC was less than two. Models were limited to three predictor variable to minimise overfitting. Since GAMMs can account for data that are not normally distributed, models were fitted with untransformed data using a Poisson distribution. Once the final model had been decided, the model residuals were assessed for heterogeneity and overdispersion. If a model was considered overdispersed, the process was repeated but this time using the negative binomial distribution. Models with a negative binomial distribution were also assessed using a forward stepwise selection of the k value. All GAMM analyses were performed using the ‘GAMM4’ package in R [61].

## Results

### Summary of baited remote underwater video deployments

A total of 107 stereo-BRUVs were successfully completed, with 64 deployments on the shallow reef and 43 deployments on the mesophotic reef (Table 2). A total of 7368 individuals (sum of MaxN) from 96 species, representing 53 families were recorded (Table 2 and Supplementary Material). A total of 79 species were recorded on shallow reef, of which 49 species were unique (Table 2). A total of 47 species were recorded on mesophotic reef, of which 17 species were unique (Table 2). Thirty species were found to occur on both shallow and mesophotic reef (Table 1).

**Table 2.** A summary of the number of BRUVS and species compositions recorded from BRUVS deployed on shallow and mesophotic reef.

|  | Shallow (20-40 m) | Mesophotic (80-110 m) |
| --- | --- | --- |
| No. of BRUV deployments | 64 | 43 |
| Species richness (SR) | 79 | 47 |
| Mean SR (± SE) per BRUV | 19(0.48) | 9(0.47) |
| Family richness | 42 | 35 |
| No. rare species | 26 | 17 |
| No. of species unique to reef depth | 49 | 17 |

Rare species were seen on fewer than three occasions.

Labridae and Monacanthidae were the most speciose families with nine species each, equating to 19 % of the total species richness. On shallow reefs, *Ophthalmolepis lineolatus* was the most ubiquitous species, being recorded on 100 % of deployments, followed by *Notolabrus gymnogenis* (94 %) and *Chrysophrys auratus* (92 %). In comparison, on mesophotic reefs *Centroberyx affinis* was the most ubiquitous species, being recorded on 74% of deployments, followed *by Nemodactylus douglasii* at 72 % and *Trachurus novaezelandiae* at 60 %. Two threatened species, *Epinephelus daemelii* (black cod) were recorded on shallow reef and *Carcharias taurus* (grey nurse shark) were recorded on shallow and mesophotic reef.

### Fish assemblage spatial distribution in relation to environment

The best fitting RDA model to describe the transformed species assemblage data included the factors depth, latitude, fished, habitat (Adj. R^2^ = 0.27, F = 5.82, P <0.001). Permutation tests of each of these constraints gave significant marginal terms (depth: *F* = 30.35 *P* <0.01, latitude: *F* = 4.01 *P* <0.01, fished: *F* = 2.24 *P* = 0.02, habitat: *F* = 1.99 *P* < 0.01). The reef metrics, relief, rugosity, ruggedness, curvature and slope were not significant in terms of explaining the transformed species assemblage data. The RDA ordination showed a clear division in BRUV deployments on shallow reefs and mesophotic reefs (Fig. 2). The majority of shallow BRUV deployment had a positive RDA1 values, while the majority of BRUV deployments on mesophotic reefs had negative RDA1 values (Fig 2). The RDA2 axis was mainly driven by habitat and latitude (Fig. 2). Mesophotic reefs were characterised by the schooling species *Trachurus novaezelandiae*, *Centroberyx affinis* and *Meuschenia scaber*, while shallow reefs were characterised by the schooling species *Atypichthys strigatus* and *Scorpis lineolata,* as well the indicator species *Chrysophrys auratus* and *Ophthalmolepis lineolatus* (Fig. 2).

**Fig. 2.**
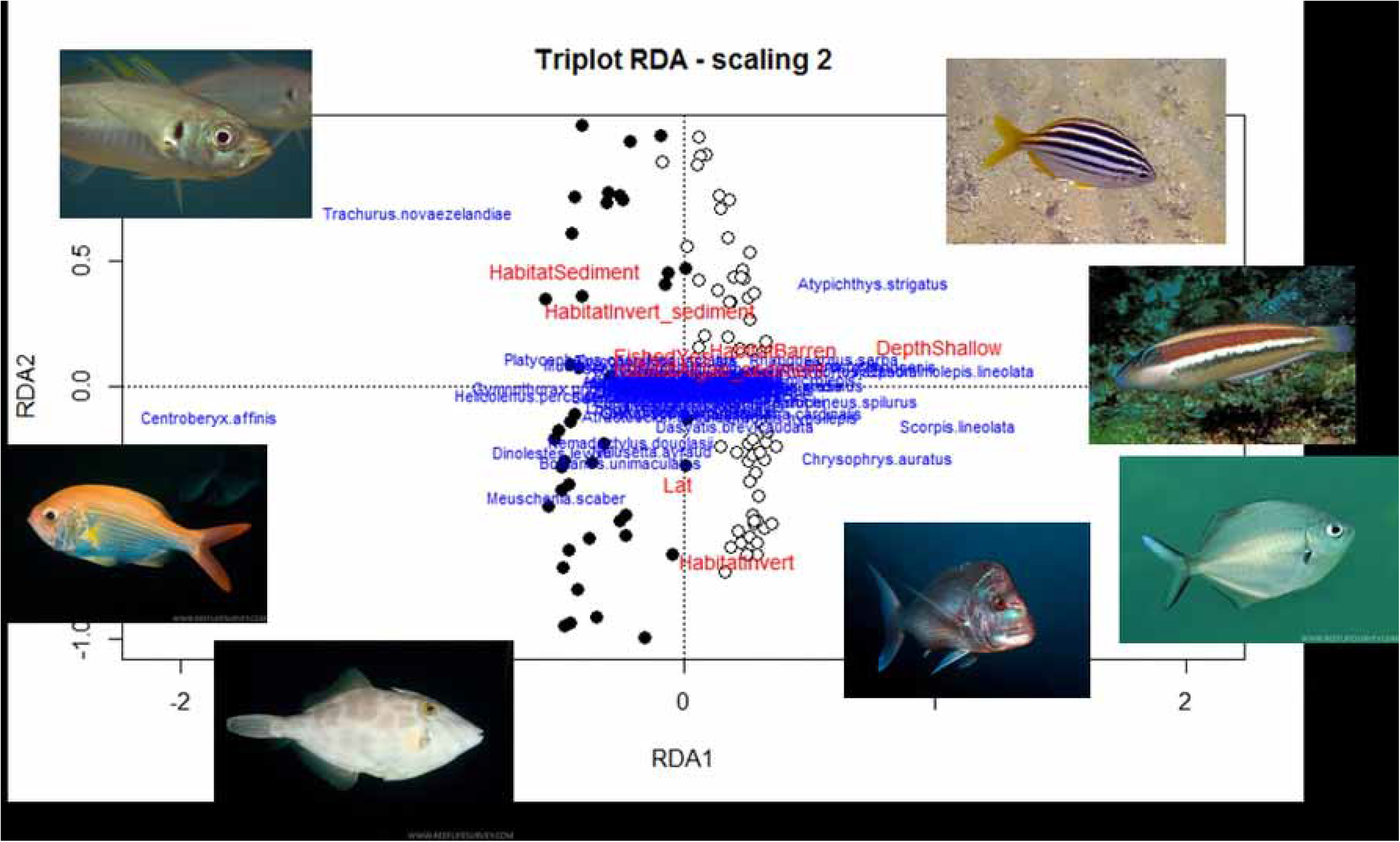
An RDA triplot ordination of transformed relative abundance data constrained by depth, latitude, fished/no-take. Filled circles represent mesophotic reef BRUV deployments and open circles represent shallow reef BRUV deployments.

### Species richness, relative abundance and family spatial distribution

The species richness recorded on shallow reef BRUVs was nearly double that recorded on mesophotic reef BRUVs, with little variation between deployments (Fig. 3). The most parsimonious generalised additive mixed model (GAMM) for species richness included the factors depth, substrate and rugosity (Table 3). BRUVs that landed on top of rocky reef had the highest species richness, with a significant positive relationship with rugosity. The total relative abundance (total MaxN) of all fishes followed a similar pattern, with more than double the number of fishes recorded on shallow reefs compared with mesophotic reefs (Fig. 3). Depth and habitat provided the most parsimonious GAMM model (Table 3). Sites that were dominated by urchin barrens and sediment habitats had the highest total MaxN. This pattern is driven by the high numbers of schooling species of fish such as *Atypichthys strigatus* and *Trachurus novaezelandiae* that commonly occurred across the shallow reefs.

**Table 3.**
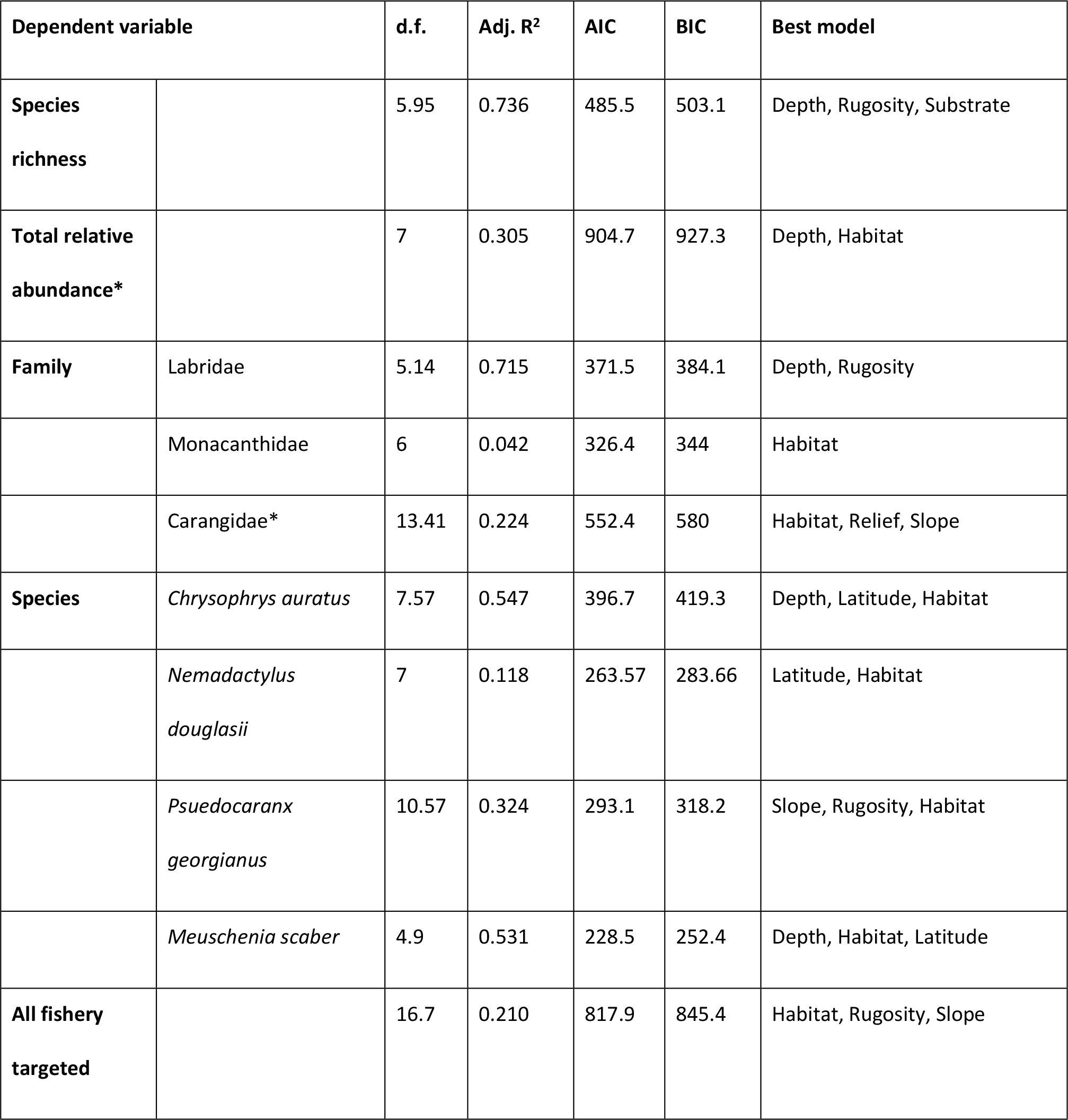
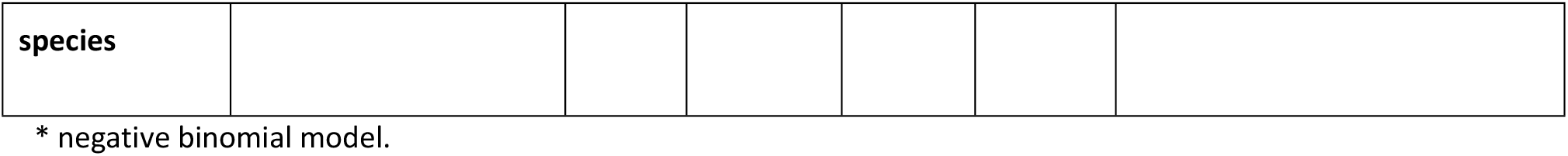
The best generalized additive mixed models (GAMMs) for predicting, species richness, relative abundance of key families, relative abundance of most targeted species and all targeted species pooled.

**Fig. 3.**
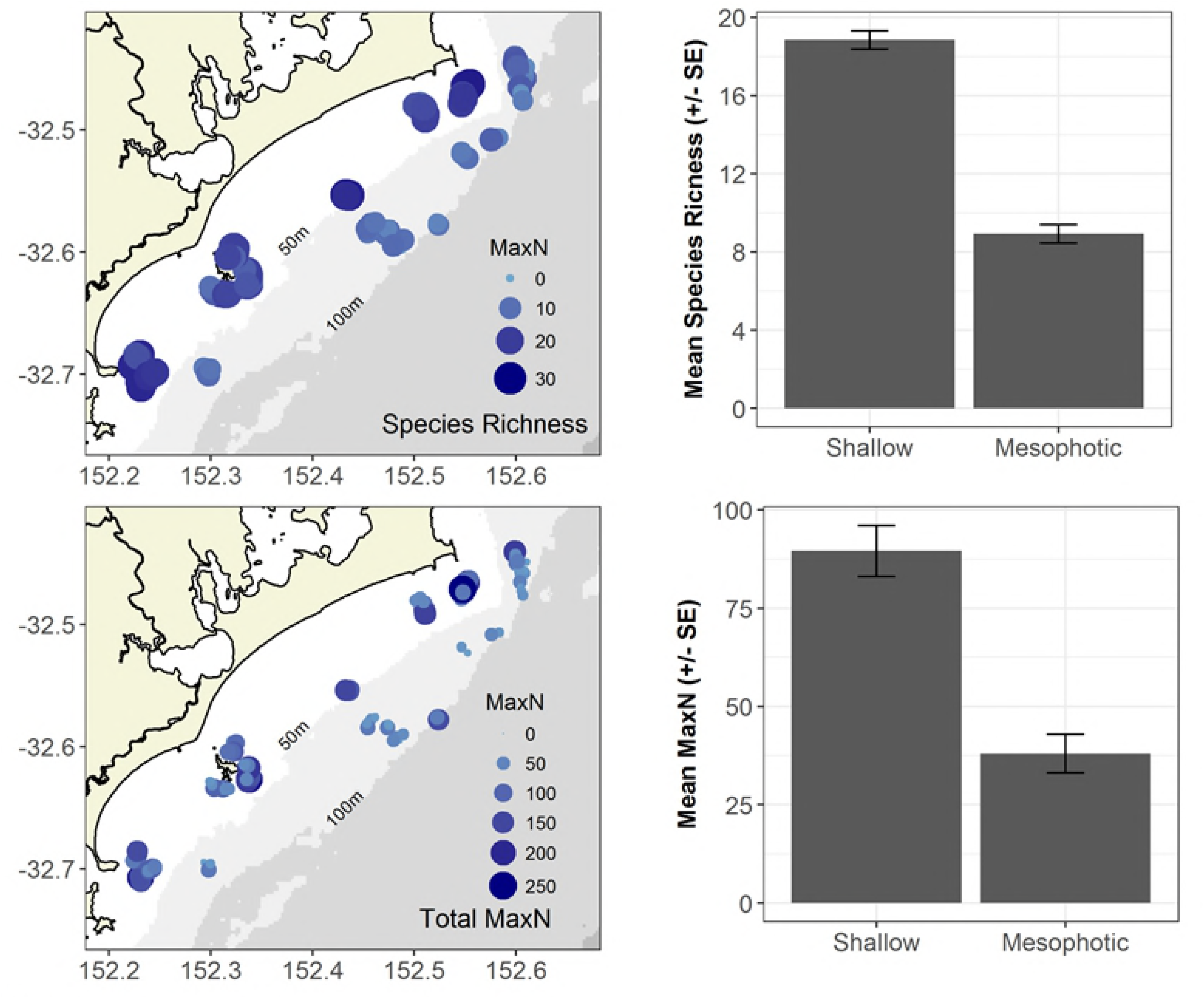
The spatial distribution of species richness and total MaxN. Top left: Distribution of species richness as observed by stereo-BRUVs across the study area. Bubble size and colour represents the species richness for each individual stereo-BRUV deployment. Top right: Mean (+/- SE) species richness across shallow and mesophotic reef. Bottom left: Distribution of total MaxN as observed by stereo-BRUVs across the study area. Bubble size and colour represents the total MaxN for each individual stereo-BRUV deployment. Top right: Mean (+/- SE) total MaxN across shallow and mesophotic reef.

The family Labridae was much more abundant on shallow reefs compared to mesophotic reef (Fig. 4). All nine species of Labridae were recorded on shallow reefs, with *Ophthalmolepis lineolatus* and *Coris Picta* being the most relatively abundant. Only one species of Labridae, *Bodianus unimaculatus*, was recorded on mesophotic reefs. The family Monacanthidae were more equally distributed across reef type (Fig. 4), with all nine species occurring on the shallow reefs, while three species were recorded on both reef types. *Meuschenia scaber* was the most abundant species on both reef types. Monacanthidae were evenly distributed across both shallow and mesophotic reefs, and the best model included the single factor habitat (Fig 4, Table 3). Habitats dominated by a high abundance of sessile invertebrates had the highest relative abundance of Monacanthids.

**Fig. 4.**
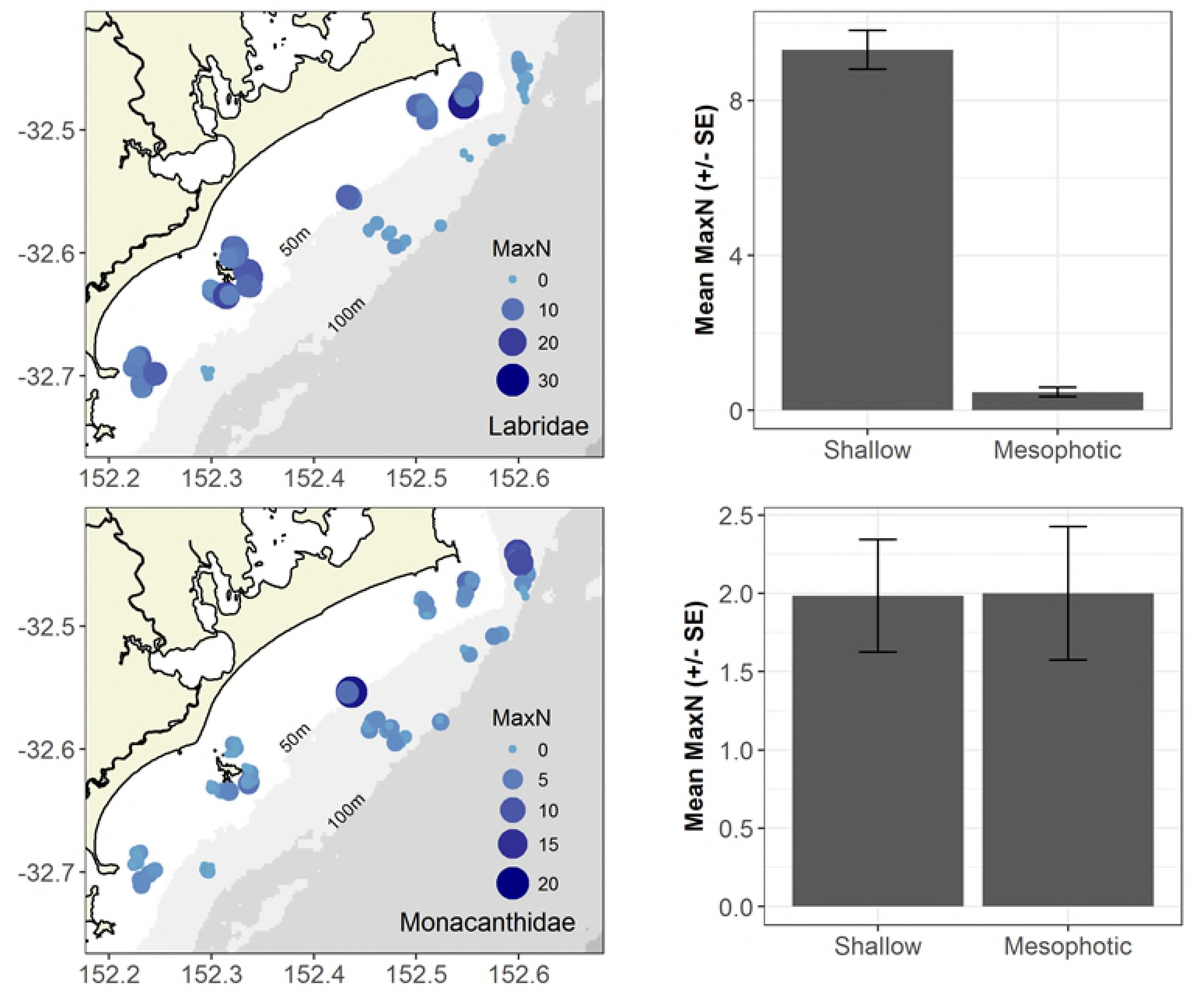
The spatial distribution of the most speciose families. Top left: Distribution of the family Labridae as observed by stereo-BRUVs across the study area. Bubble size and colour represents the total MaxN of the family Labridae for each individual stereo-BRUV deployment. Top right: Mean (+/- SE) total MaxN for the family Labridae across shallow and mesophotic reef. Bottom left: Distribution of the family Monacanthidae as observed by stereo-BRUVs across the study area. Bubble size and colour represents the total MaxN of the family Monacanthidae for each individual stereo-BRUV deployment. Top right: Mean (+/- SE) total MaxN for the family Monacanthidae across shallow and mesophotic reef.

The distribution of fishery targeted species across the two depth categories was highly variable and species dependent. The spatial distribution of the most targeted species (*Chrysophrys auratus*) was related to depth, latitude and habitat type (Table 3). On average, the relative abundance of *C. auratus* on shallow reefs was six times greater than on mesophotic reefs (Fig. 4), although, there was greater variability between BRUV deployments on shallow reef (Fig. 4). The positive effect of latitude showed that abundances of *C. auratus* were highest at the Seal Rocks sites; the most northern survey site (Fig 5). Also, *C. auratus* occurred in greater abundance on BRUV deployments that were located on the edge of reefs that were dominated by invertebrate/sediment or sediment habitats. *Nemadactylus douglasii*, also a highly targeted species, was more evenly distributed across reef type, with latitude and habitat type providing the best model to describe the spatial distribution of this species (Table 3). The positive relationship between latitude and relative abundance of *N. douglasii* showed greater abundance at Seal Rocks sites (Fig. 5). The spatial distribution of the carangid *Psuedocaranx georgianus* was far more variable than *C. auratus* and *N. douglasii*. However, on average the relative distribution was similar across the two depth categories (Fig. 5). *Psuedocaranx georgianus* tend to be observed in small schools on low rugosity reef edges. The most parsimonious model that best described the spatial distribution of *P. georgianus* included the factors slope, rugosity and habitat (Table 3). The spatial distribution of the monacanthid *Meuschenia scaber* was highly variable, but on average higher relative abundances was observed on mesophotic reefs (Fig. 5). The most parsimonious model best described the spatial distribution of *M. scaber* included the factors depth, habitat and latitude (Table 3). It was sites within the mid latitudes of this study that had the highest relative abundances and the higher relative abundances tended to be on low relief reef (Fig. 5).

**Fig. 5.**
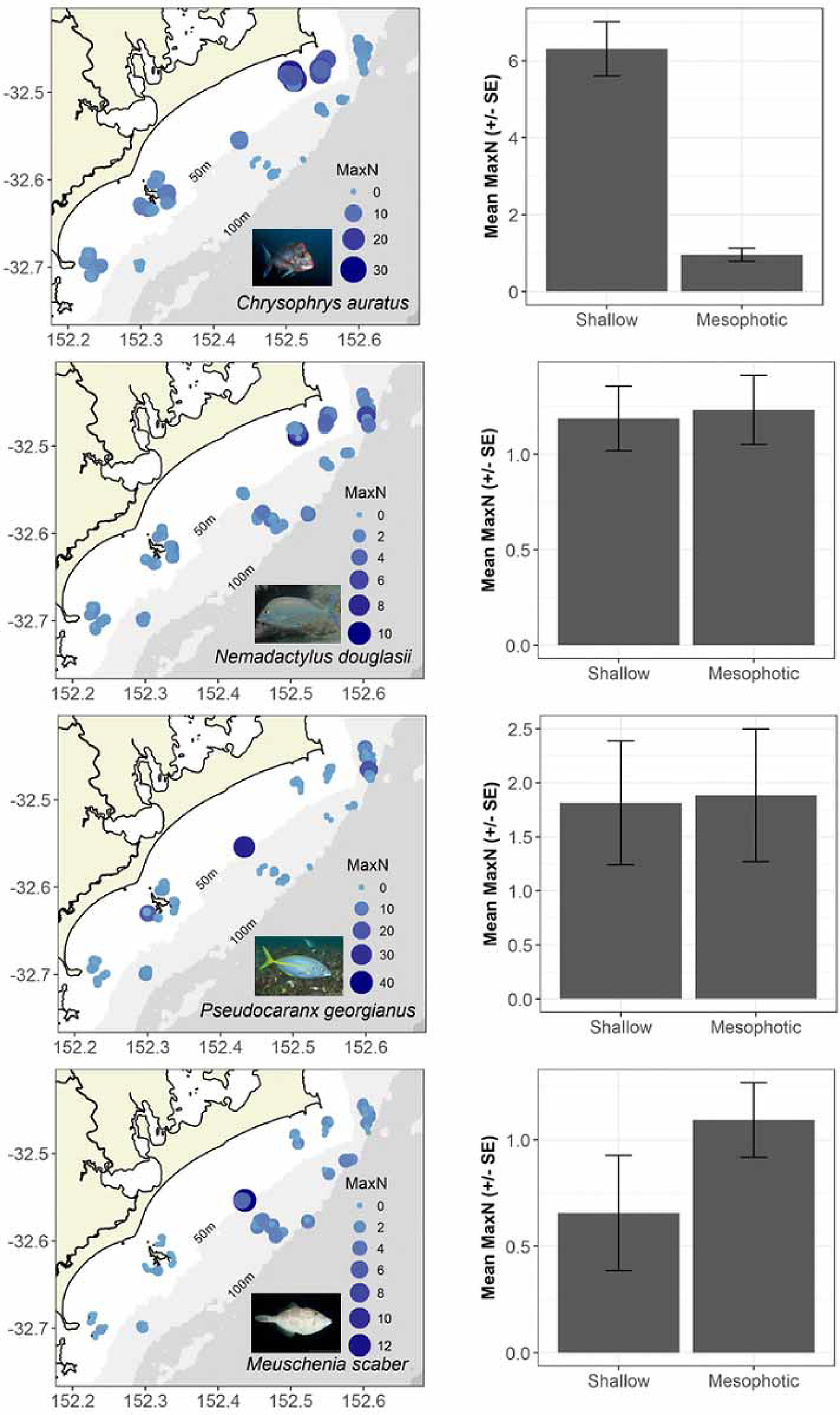
Spatial distribution of species of interest. Top row left: Distribution of the *Chrysophrys auratus* as observed by stereo-BRUVs across the study area. Bubble size and colour represents the MaxN of *Chrysophrys auratus* for each individual stereo-BRUV deployment. Top row right: Mean (+/- SE) MaxN of *Chrysophrys auratus* across shallow and mesophotic reef. Second row left: Distribution of the *Nemadactylus douglasii* as observed by stereo-BRUVs across the study area. Bubble size and colour represents the MaxN of *Nemadactylus douglasii* for each individual stereo-BRUV deployment. Second row right: Mean (+/- SE) MaxN of *Nemadactylus douglasii* across shallow and mesophotic reef. Third row left: Distribution of the *Psuedocaranx georgianus* as observed by stereo-BRUVs across the study area. Bubble size and colour represents the MaxN of *Psuedocaranx georgianus* for each individual stereo-BRUV deployment. Third row right: Mean (+/- SE) MaxN of *Psuedocaranx georgianus* across shallow and mesophotic reef. Bottom row left: Distribution of the *Meuschenia scaber* as observed by stereo-BRUVs across the study area. Bubble size and colour represents the MaxN of *Meuschenia scaber* for each individual stereo-BRUV deployment. Bottom row right: Mean (+/- SE) MaxN of *Meuschenia scaber* across shallow and mesophotic reef.

Species that are actively targeted and highly retained by both recreational and commercial fishers showed a relatively equal distribution across both shallow and mesophotic reefs (Fig. 6). Habitat, rugosity and slope best described the variability between sites (Table 3). Reef dominated by algae and reef edge habitats had the highest abundance of fishery-targeted species. While there was a strong positive relationship between fishery-targeted species and reef rugosity, there was a weak negative relationship with slope.

**Fig. 6.**
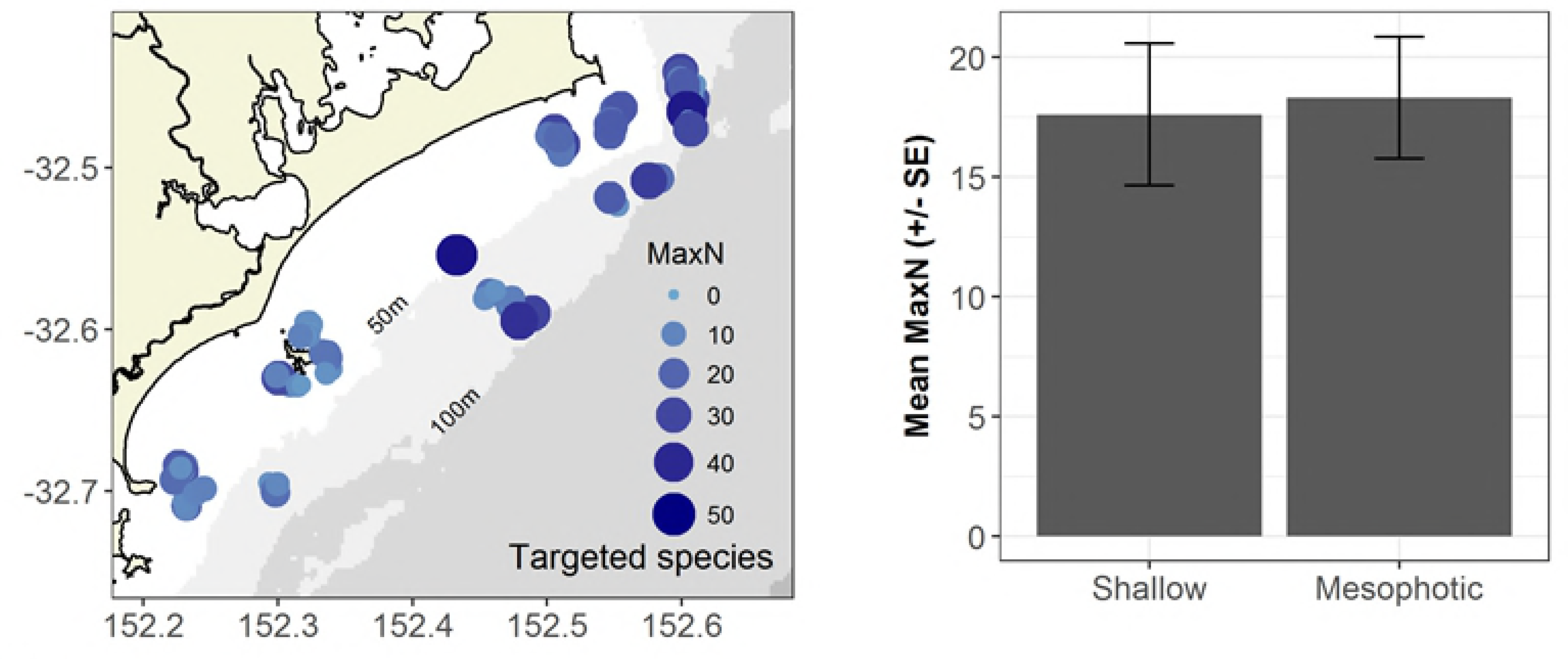
The spatial distribution of fishery targeted species. Left: Distribution of all commercial and recreationally targeted species as observed by stereo-BRUVs across the study area. Bubble size and colour represents the MaxN of all commercial and recreationally targeted species for each individual stereo-BRUV deployment. Top row right: Mean (+/- SE) MaxN of all commercial and recreationally targeted species across shallow and mesophotic reef.

On average, the lengths of *C. auratus* on mesophotic reefs were larger than on shallow reef, with the mean slightly below, and upper confidence intervals above, the minimum legal length for retention of this species (Fig. 7). Mesophotic reefs also had a greater proportion of legally sized fish at 62 % compared to shallow reefs 12 % (Fig 7). In comparison, the mean lengths and proportion of fish above the MML of *N. douglassi* were very similar between mesophotic and shallow reef (Fig. 7).

**Fig. 7.**
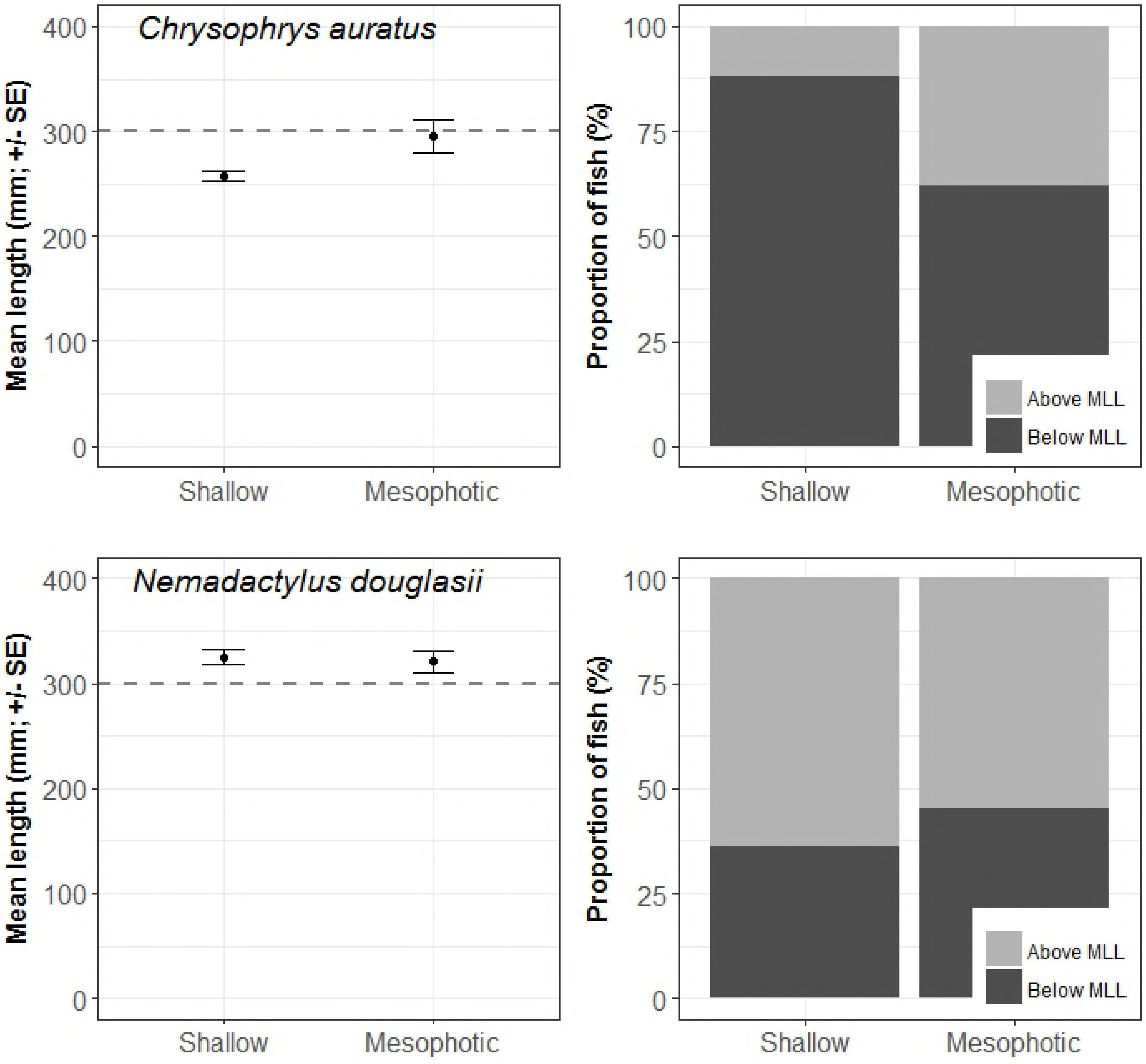
Lengths of two of the prominent fishery targeted species. Left: The mean total lengths of Chrysophrys auratus and Nemadactylus douglasii with the minimum legal length (MML) marked as dashed line. Right: The proportion of measured fish above and below the MML.

## Discussion

The fish assemblages on rocky reefs at mesophotic depths (80-110m) were found to be distinct from those associated with shallow rocky reefs (20-40m). Despite the large differences in species richness and relative abundance of all fishes, 30 species (i.e. 31 % all species recorded during this study) occurred across the depth gradient sampled. This is the first study to compare rocky reef fish assemblages across a 80-90 m depth gradient at these latitudes, and there are few comparable temperate studies. The vast majority of mesophotic reef research has occurred in tropical systems in Australia and USA / central America [13]. There are even fewer studies (∼1 %) using BRUVs, particularly stereo-BRUVs, to sample fish and habitats at mesophotic depths [13]. The species richness that we recorded from stereo-BRUVs on these temperate mesophotic reefs is approximately half of what has been recorded using BRUVs in tropical systems [4,33,45]. However, the species richness to depth gradient relationship is consistent across both tropical and temperate systems [Current study, 4,62].

The transition zone between the shallow and mesophotic species assemblages is unclear in this study as we did not sample between depths of 40-80 m. The mesophotic reef that was sampled during this study is disconnected from rocky reefs found in shallow waters as it is separated by large expanses of soft sediment habitats [54]. The current referenced global definition for the transition to mesophotic ecosystems is 40 m [11,63], but, this will be regionally specific and dependent on factors such as light, temperature and habitat [13]. The connectivity across the depth gradient is also a significant unknown. An understanding of depth connectivity is needed to determine if mesophotic reefs can provide refuge from short term pressures such as storms or heatwaves or long term pressures such as fishing. There is evidence that mesophotic reefs can provide ‘refuge’ habitat for fishery targeted species [33–35,64]. Some caution is needed when making generalisations of connectivity across depths ranges, however, as there is evidence of differing life histories and intraspecific variability in demographic traits of fishes using reefs at mesophotic depths [65].

This study found strong depth related patterns that could be coupled with depth related reef and habitat complexity [4,6,20]. In all but two models (Species richness and Labridae), the factor habitat was selected, highlighting the importance of habitat in describing the distribution of fishes. The correlation between increasing depth and decreasing light equates to a change in habitat structure, with the shallower reefs dominated by macroalgae [66,67], and at depths >30m sessile invertebrates, such as, sponges and octocorals. As the shallow reefs surveyed in this study were in 20-40 m depth range, a transition between algae and sessile invertebrate dominated habitats was observed. Therefore, herbivorous and omnivorous fishes are more likely to inhabit shallow reefs where algae is present and thus increasing species richness and abundance. In comparison, at mesophotic depths you would expect to only observe the carnivorous, planktivorous, and scavenger fish and species with greater tolerances to ocean currents and thermoclines.

Apart from light availability and habitat, ocean currents and temperature gradients or thermoclines further separate mesophotic reefs from adjacent shallow reefs [4]. In temperate Eastern Australia, the East Australian Current (EAC) has the greatest influence on the oceanography and connectivity of deeper reefs [68,69]. This is particularly relevant as the strength and the seasonality of the EAC has recently changed with warmer water pushing further south and for longer periods of time [70]. As the EAC is at its strongest (fastest flowing and warmer temperatures) across these mesophotic reefs, the EAC has the potential to influence range extension or a change in the distribution of fishes. Therefore, the EAC has the potential to influence biodiversity, abundance of fishery targeted species and species of conservation significance [69]. Further seasonal sampling is required to test hypotheses about the effects of seasonality and the EAC, i.e. differences between warm and cold water periods.

The physical structure of the reefs (rugosity, slope and relief) is likely to have the greatest influence on the spatial distribution of fishes on rocky reefs. The swath acoustic data was beneficial in selecting reefs to sample, but it can also be used to derive metrics that can possibly predict the spatial distribution of species richness, species of interest and all fishery target species pooled together. The derivation of habitat metrics from swath acoustic data can provide various levels of explanatory or predictive ability relating to fish assemblage composition and distribution [21,25,28,71,72]. In this study rugosity, slope or relief were selected in the ‘best’ models explaining the variability in many aspects of the fish assemblages that were sampled. While many studies focus on explaining or predicting the spatial distribution of individual species, a more relevant application for spatial planning would be to use reef metrics to explain or predict at higher levels such as species richness or pooled fishery targeted species. For this study, a combination of depth, substrate type, habitat type and rugosity best described species richness and fishery targeted species. These two variables could provide managers with information on areas of high fish diversity or fishery significance.

At a species level, the explanatory factors varied among the four species that were selected a priori for analysis. *Chrysophrys auratus* is arguably the most important recreational and commercial fishery in this region [48]. While *C. auratus* were twice as abundant on shallow reef, on average *C. auratus* were larger around mesophotic reefs. In proportion, there were more *C. auratus* above the minimum legal length for retention and thus considered sexually mature [73] on mesophotic reefs. This supports the hypothesis that shallow inshore reefs provide important habitat for juvenile *C. auratus,* while deeper mesophotic reef provide additional habitat for *C. auratus*. A recent hypothesis is that fishing pressure drives ontogenetic ‘deepening’ in exploited species [74]. *Chrysophrys auratus* is known to have a relatively small home range, but in many regions they can move hundreds of kilometres [75–77]. It has been demonstrated that *C. auratus* have small home ranges and display strong site fidelity on shallow reefs in the PSGLMP [75], but their movements and use of deeper reef habitats is unknown and warrants further investigation.

In contrast, the distribution of *Nemadactylus douglassi* was more influenced by latitude than by depth. They do occur across a range of habitats including soft sediment and rocky reef and mainly feed on soft sediment associated molluscs and crustaceans [78,79]. *Nemadactylus douglassi* are medium to large bodied fish that are targeted by recreational and commercial fishers. On average, the *N. douglassi* recorded during this study were above the minimum legal length for retention and could be considered sexually mature [79]. On this section of coastline the relative abundance and lengths of *N. douglassi* is fairly constant across the depth gradient. Similarly, *Psuedocaranx dentex* was commonly observed on both types of reef, but they were patchier in their distribution. They were often observed in large schools on BRUV deployments positioned on the edge of reef or over adjacent soft-sediments. They too are targeted and retained by recreational and commercial fishers [79]. *Meuschenia scaber* is the most numerically abundant Monacanthid species in eastern Australia and New Zealand, but very little is known of its ecology and biology [80]. They are known to inhabit a wide range of depths but in this study there appears to be a preference for deeper mesophotic reefs. Feeding exclusively on sessile invertebrates such as sponges, ascidians, polyzoans, hydroids and barnacles, *M. scaber* is well suited to these mesophotic reefs [80]. For *N. douglassi* and *P. dentex* there doesn’t appear to be any preference for a particular depth of reef and nor is there any clear ecological rationale for one.

This study demonstrated that the fish assemblages of rocky reef at mesophotic depths are statistically different to the adjacent shallow reef systems. Despite more than double the total abundances, there were similar relative abundances of fishery target species across both shallow and mesophotic reefs suggesting that, from a fisheries management perspective, these reef systems have the potential for similar social and economic values. Historically, mesophotic reefs were fished more by commercial fishers than by recreational anglers. However, increased knowledge and access to improved technology is now allowing boat-based recreational fishers to target deeper reefs. Swath acoustic data contributed to explaining the spatial distribution of each aspect of the fish assemblage. There is still a research need to investigate seasonal patterns and fine-scale intra-reef variability in fish assemblages on these temperate mesophotic reefs. The use of a complementary method, such as ROV or towed video that passively samples the fish assemblages would provide valuable information on the species not captured through the use of baited remote underwater video sampling [81]. Notwithstanding this limitation, this study has clearly demonstrated that reefs at mesophotic depths are important and should be taken more into consideration by managers when zoning marine protected areas.

## Acknowledgements

The study was conducted with the support of the NSW Department of Primary Industries and the NSW Office of Environment and Heritage. We would like to thank Roger Laird and Tom Davis for their assistance with the field work. Thank you to Bob Creese who provided valuable feedback and revisions. Photos are courtesy of David Harasti, Reef Life Survey and Fishes of Australia.

